# Ionic strength and calcium regulate the membrane interactions of myelin basic protein and the cytoplasmic domain of myelin protein zero

**DOI:** 10.1101/529586

**Authors:** Arne Raasakka, Nykola C. Jones, Søren Vrønning Hoffmann, Petri Kursula

## Abstract

The formation of a mature myelin sheath in the vertebrate nervous system requires specific protein-membrane interactions. Several myelin-specific proteins are involved in the stacking of lipid membranes into multilayered structures around neuronal axons, and misregulation of these processes may contribute to chronic demyelinating diseases. Two key proteins functioning in myelin membrane binding and stacking are the myelin basic protein (MBP) and protein zero (P0). Other factors, including Ca^2+^, are important for the regulation of myelination. Here, we studied the effects of ionic strength and Ca^2+^ on the direct molecular membrane interactions of MBP and the cytoplasmic domain of P0 (P0ct). While both MBP and P0ct bound and aggregated negatively charged lipid vesicles, while simultaneously folding, both ionic strength and calcium had systematic effects on these interactions. Especially when decreasing membrane net negative charge, the level and kinetics of vesicle aggregation, which is a functional assay for myelin membrane-stacking proteins, were affected by both salt and Ca^2+^. The results indicate that the effects on lipid membrane surfaces by ions can directly affect myelin protein-membrane interactions at the molecular level, in addition to signalling effects in myelinating glia.

## Introduction

Myelin contributes greatly to the efficiency of both the central and peripheral nervous system (CNS and PNS respectively) *via* axonal insulation and trophic support (1, 2). Comprised of tightly stacked lipid membranes, held together by a specific assortment of proteins, compact myelin (CM) – a water-deficient structure – insulates selected axonal segments and enables fast saltatory nerve impulse conduction. While neuronal axons undergo frequent membrane depolarization, and [K^+^] and [Na^+^] vary upon action potential firing, the ionic conditions in myelinating glia remain fairly constant, with higher [K^+^] and [Na^+^] compared to neurons in general (3).

In myelinating cells, [Ca^2+^] and [Zn^2+^] (1 mM and 50 µM, respectively) are relatively high compared to many other cell types (3, 4). This signifies the importance of the underlying Ca^2+^ signalling (5-7); due to the narrow bilayer spacing and low water content in CM, intracellular divalent cations must be subjects of attractive and repulsive interactions. These involve negatively charged and zwitterionic phospholipid headgroups, as well as myelin proteins, which are often highly positively charged and directly participate in interactions with other proteins and lipids. Both Ca^2+^ and Zn^2+^ have been studied with respect to autoinflammatory diseases and pathogenesis (5, 8-10), and intracellular Ca^2+^ levels and changes therein have been directly linked to the correct development of myelin (11-14).

Myelin basic protein (MBP) plays a fundamental role in forming and maintaining stable myelin membrane stacks, defining the boundaries between CM and non-compact myelin (15-18). MBP is involved in multiple sclerosis (MS) (19-21), being an intrinsically disordered protein (IDP) that folds upon lipid binding-induced charge neutralization 16, 22, 23). MBP is highly positively charged and manifests as a pool of post-translationally citrullinated variants with differing net charges (24). MBP requires negatively charged lipids for membrane adhesion, but correct myelin morphology is also dependent on the presence of other lipids and a controlled ionic content (14, 16, 25). MBP interacts with divalent cations (26-30), with the underlying mechanisms being very specific; one would expect electrostatic repulsion due to its unusually high positive net charge. MBP binds to calmodulin in a Ca^2+^-dependent manner (27, 31-33), and the pool of different MBP isoforms has been shown to regulate oligodendroglial Ca^2+^ influx (34, 35). Concentration-dependent binding of divalent cations to different MBP variants could, together with altered membrane compositions, influence their *in vivo* functions, having implications in MS etiology and the stability of CM (6, 14, 27, 36).

Myelin protein zero (P0) is a transmembrane protein in PNS myelin, with relevance in human peripheral neuropathies and animal models (37-39). It consists of an extracellular domain with an immunoglobulin-like fold (40), a single transmembrane helix, and a short cytoplasmic segment (P0ct) involved in the regulation of membrane fluidity (41). Similarly to MBP, P0ct is highly positively charged and directly interacts with phospholipids through electrostatics, gaining secondary structure in the process (41, 42), which might make it susceptible to effects caused by cationic species.

In this study, the effects of ionic strength and Ca^2+^ on the structural and functional properties of recombinant MBP and P0ct binding to lipid membranes were studied using different biophysical techniques. The results shed new light on the interplay between ions and proteins in CM and support the proposed role of Ca^2+^ as a regulator of the bilayer-MBP interaction (14, 36, 43).

## Experimental procedures

### Protein expression and purification

Mouse MBP and human P0ct were expressed and purified as described (16, 41). The final size-exclusion chromatography after affinity purification and tag removal was performed using Superdex75 16/60 HiLoad and Superdex75 increase 10/300GL columns (GE Healthcare) with 20 mM HEPES, 150 mM NaCl, pH 7.5 (HBS) as mobile phase. When required, the buffer was exchanged to water by sequential dialysis. The purified proteins were either used fresh or snap-frozen in liquid N_2_ and stored at −80 °C for later use.

### Vesicle preparation

1,2-dimyristoyl*-sn*-glycero-3-phosphocholine (DMPC), 1,2-dimyristoyl*-sn*-glycero-3-phospho-(1’-rac-glycerol) (Na-salt) (DMPG), and the deuterated d_54_-DMPC, d_54_-DMPG, and d_7_-cholesterol were from Avanti Polar Lipids (Alabaster, Alabama, USA). The detergent *n*-dodecylphosphocholine (DPC) was from Affymetrix.

Lipid stocks were prepared by dissolving the dry lipids in chloroform or chloroform:methanol (9:1 v/v) at 10 – 30 mM. All mixtures were prepared from the stocks at desired molar ratios, followed by solvent evaporation under an N_2_ stream and freeze-drying overnight at −52 °C under vacuum. The dried lipids were either stored air-tight at −20 °C or used fresh for liposome preparation.

Liposomes were prepared by agitating dried lipids with either water or HBS at a concentration 10 – 15 mM, followed by gentle mixing at ambient temperature, to ensure that no unsuspended lipids remained in the vessel. MLVs were prepared by freeze-thawing using liquid N_2_ and a warm water bath, with vigorous vortexing afterwards. Such a cycle was performed 7 times. SUVs were prepared by sonicating fresh MLVs using a probe tip sonicator (Materials Inc. Vibra-Cell VC-130) until clear, while avoiding overheating. All lipid preparations were immediately used for experiments.

### Vesicle turbidimetry

For turbidimetric measurements, samples containing 0.5 mM DMPC:DMPG (1:1) SUVs were prepared with 2.5 µM P0ct or MBP in duplicate at 150 µl final volume on a Greiner 655161 96-well plate. Optical density at 450 nm was recorded immediately after thorough mixing at 25 °C using a Tecan Spark 20M plate reader.

### Synchrotron radiation circular dichroism spectroscopy

SRCD data were collected from 0.1–0.4 mg ml^−1^ protein samples in water or 0.5% SDS on the AU-CD beamline at ASTRID2 (ISA, Aarhus, Denmark). Samples containing lipids were prepared immediately before measurement by mixing P0ct (P/L ratio 1:200) or MBP (P/L ratio 1:300) with freshly sonicated SUVs. 100-µm and 1-mm pathlength closed circular cells (Suprasil, Hellma Analytics) were used for the measurements, and SRCD spectra were recorded from 170 to 280 nm at 30 °C and truncated based on detector voltage levels as well as noise. Baseline spectra were subtracted from sample spectra, and CD units were converted to Δε (M^−1^ cm^−1^) using CDtoolX (44).

### Stopped-flow SRCD

Rapid kinetic SRCD data were collected using an SX-20 stopped-flow instrument (Applied Photophysics) mounted on the AU-AMO beamline at ASTRID2 (ISA, Aarhus, Denmark) at 30 °C. Acquisition of kinetic data comprised of 6–10 shots (160-µl total volume per shot) per sample. The two 1-ml syringes were loaded as follows: syringe 1 contained 0.1 mg/ml protein and syringe 2 a lipid solution (final molar P/L ratios of 1:200 and 1:300 for P0ct and MBP, respectively). The contents per each shot were rapidly mixed (2 ms dead time) before injection into the measurement cell (2-mm pathlength). Change in CD signal (mdeg) was monitored at 195 nm for 5 s per shot. The successful overlaying shots were averaged into a single curve for each sample. All sample sets were prepared and measured in duplicate. Water baselines were subtracted from sample curves, and data were analyzed in GraphPad Prism 7 using single- and two-phase exponential decay functions to obtain rate constants.

## Results

In the current project, the folding and lipid membrane aggregation of MBP and P0ct were studied. Optical density measurements were used to observe the aggregation of lipid vesicles in the presence of proteins and different ionic species. The studies were expanded to different ratios of DMPC:DMPG, which affects the surface net charge of the SUVs. Using SRCD spectroscopy, the folding of the two proteins was probed, and time-resolved experiments provided data on membrane aggregation kinetics.

### The effect of ionic content on myelin protein-induced vesicle turbidity

As a functional assay for MBP and P0ct, turbidimetric experiments on SUVs were carried out. In our earlier studies, we concluded that in the presence of MBP, net negatively charged lipid vesicles aggregate and subsequently form myelin-like bilayer stacks, resulting in sample turbidity (16). P0ct, on the other hand, can cause SUVs to undergo fusion events to produce large aggregated lipid bodies (41), which may cause additional sample turbidity.

The turbidity induced by the proteins for 1:1, 4:1, and 9:1 DMPC:DMPG SUVs, at a 1:200 protein-to-lipid (P/L) ratio, was determined. The experiments were carried out at various concentrations of NaCl, NaF, and CaCl_2_. Since ionic species, especially multivalent cations, are known to induce lipid aggregation in the absence of proteins (45), control samples without proteins were included (Fig. 1). Protein-free sample turbidity was observed mostly in the presence of CaCl_2_, with more CaCl_2_ needed for induction, when the net charge of the lipids became more negative. The effect of monovalent ions as inducers of turbidity *per se* in protein-free samples was low.

**Fig 1.**
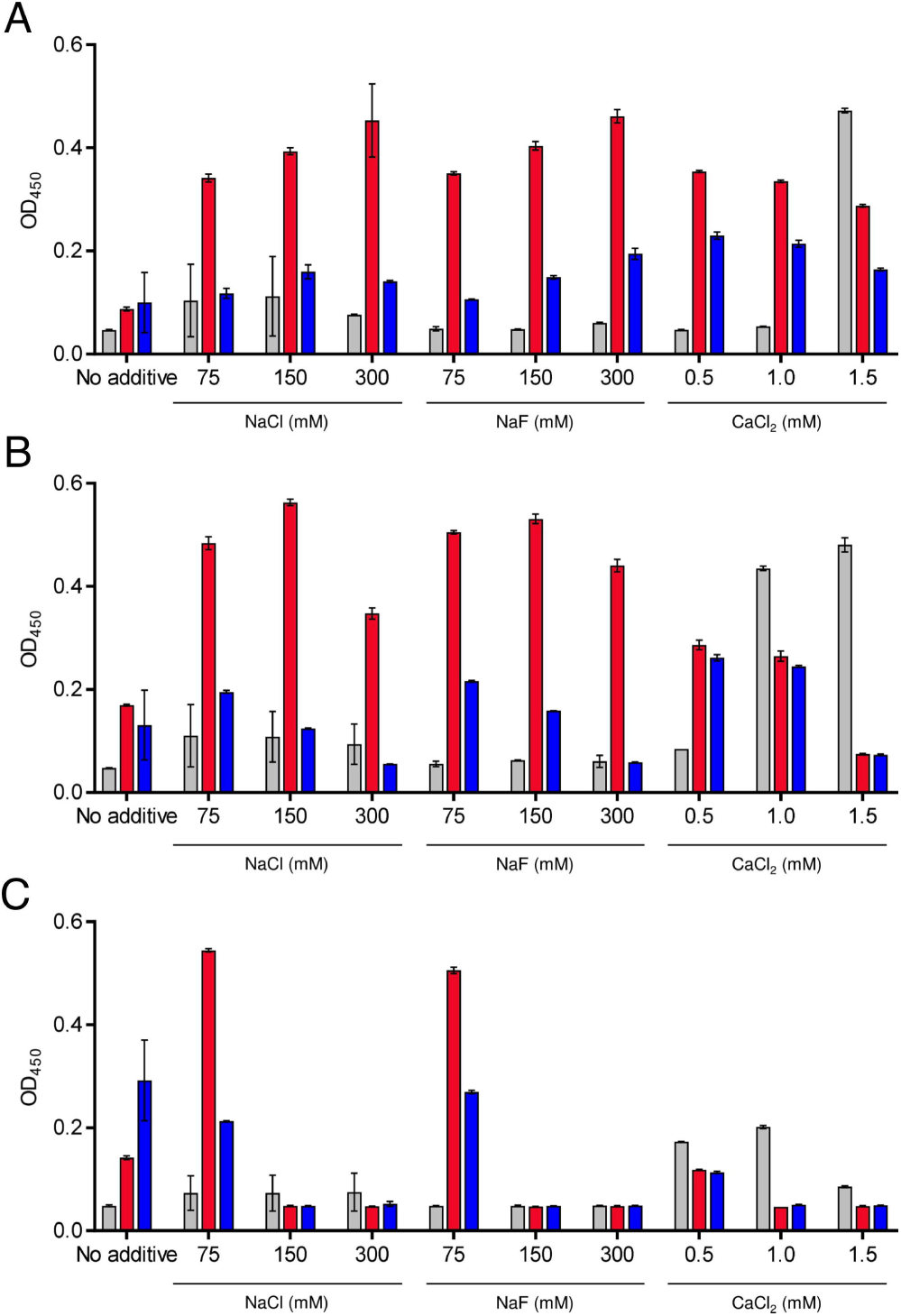
Vesicle turbidimetry. MBP (red) and P0ct (blue)-induced turbidity was studied in (A) 1:1, (B) 4:1, and (C) 9:1 DMPC:DMPG vesicles under different salt conditions. Each condition in the absence of protein is shown for reference (gray). Error bars represent standard deviation.

NaCl and NaF boosted the measured turbidity levels in all lipid mixtures at least to some extent. The turbidity induced by MBP was higher than that of P0ct, with the exception of DMPC:DMPG (9:1), in which P0ct was surprisingly more efficient than MBP. In the presence of either protein, the induced turbidity systematically increased with salt concentration DMPC:DMPG (1:1) (Fig. 1A). In DMPC:DMPG (4:1) and (9:1) the turbidity decreased above a certain ionic strength (Fig. 1B, C), indicating that the balance between ionic strength and the net charge of the lipids plays a role in protein-induced membrane aggregation. The turbidification of DMPC:DMPG (9:1) was completely abolished at ionic concentrations of 150 mM or above. In all three lipid mixtures, both proteins in the presence of NaCl and NaF produced roughly the same levels of turbidity, or at least the observed concentration-dependent trends were identical. This indicates that the choice of monovalent anion matters little in the level of induced turbidity.

In order to follow the physiological concentration regime in myelinating glia, CaCl_2_ was included at much lower concentration than NaCl and NaF, but the increased turbidity was still observed. CaCl_2_ alone was able to induce lipid turbidity in the absence of protein; the levels varied as a function of concentration. Calcium-induced turbidity showed differences between the three DMPC:DMPG ratios: the turbidification of DMPC:DMPG (1:1) vesicles was more resistant to increased CaCl_2_ concentration than 4:1 and 9:1 mixtures (Fig. 1).

CaCl_2_ changed the turbidity induced by MBP and P0ct, depending on the membrane composition. In DMPC:DMPG (1:1), turbidity was higher than in the other two lipid mixtures, being at a similar level to that induced by the presence of salt (Fig. 1A). DMPC:DMPG (9:1) showed the least turbidity, and high calcium concentrations brought OD to the baseline level (Fig. 1C). Interestingly, in DMPC:DMPG (4:1), the presence of 0.5 mM and 1 mM CaCl_2_ showed a great difference in turbidity in the absence of proteins, but when proteins were included the turbidity levels were almost identical, indicating that the proteins could override the aggregating effect of CaCl_2_ at moderate concentrations (Fig. 1B). Increasing CaCl_2_ to 1.5 mM diminished the turbidity induced by MBP or P0ct in all membrane compositions. Taking all these results together, it is evident that while salt can influence the interactions between proteins and lipids, and protein-decorated vesicles with each other, the effect of Ca^2+^ with phospholipid headgroups is rather specific. Even small changes in divalent cation levels, in the range 0.5-1.5 mM, can alter the protein-lipid interactions required for vesicle aggregation and fusion, given that negatively charged lipid species do not dominate in abundance over neutral/zwitterionic lipids.

### Folding of lipid-bound MBP and P0ct

We observed lipid-induced increase in secondary structure content for both MBP and P0ct in our earlier studies, in the absence of salt (16, 41). To investigate the effect of ionic strength and divalent cations on protein folding, SRCD was carried out with 150 mM NaF and 1 mM CaCl_2_. NaF was used due to the high UV absorbance of Cl^−^. NaF had a very similar concentration-dependent behavior to NaCl in turbidimetric studies (Fig. 1).

NaF and CaCl_2_ had no effect on MBP or P0ct overall folding in water and 0.5% SDS (Supplementary Fig. S1). The SRCD spectra acquired for proteins in the presence lipids often suffer from turbidity-induced light scattering, here, especially at 1:1 and 4:1 DMPC:DMPG ratios in the presence of CaCl_2_ (Fig. 2A, B). Under such conditions, the positions of the spectral peaks can be used to detect the presence of secondary structure. The observed changes in conformation correspond well to the turbidimetry assays under the same conditions, whereby the negative peak in SRCD at 200 nm is indicative of unfolded structure and possibly lack of membrane binding.

**Fig 2.**
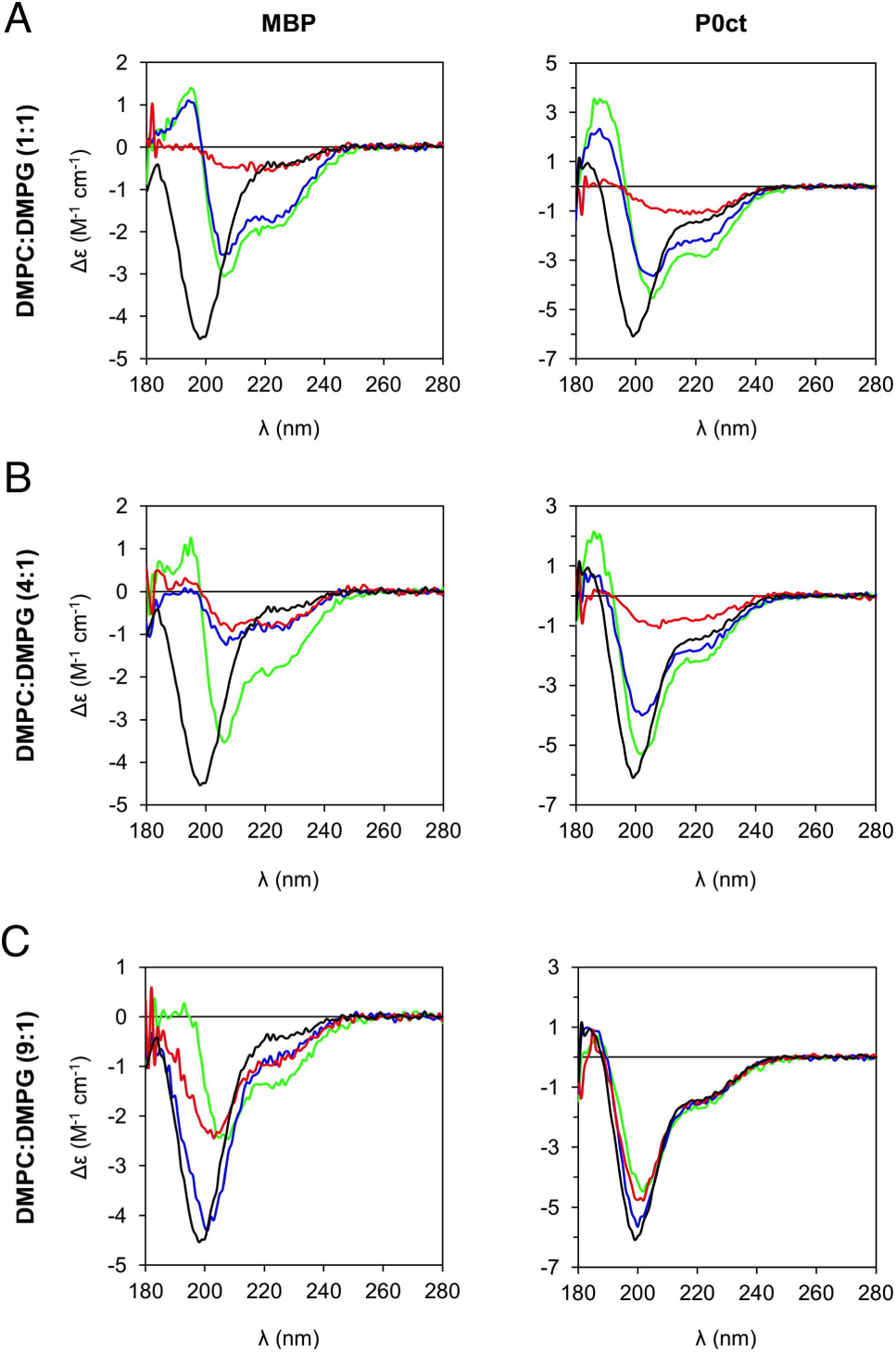
Lipid-induced protein folding in the presence of salt and divalent cations. SRCD experiments reveal differences in the folding of MBP and P0ct in different lipid compositions. Trace explanations: no additive (green); 150 mM NaF (blue); 1 mM CaCl_2_ (red). Protein controls without additives or lipids in water (black) are plotted for reference in each panel.

Without additives, MBP reached a similar secondary structure content in all lipid compositions (Fig. 2). In DMPC:DMPG (9:1), the presence of NaF or CaCl_2_ decreased MBP helical content, potentially indicating less binding to lipids (Fig. 2C). Also P0ct obtained secondary structure in lipid mixtures, and the decrease of membrane negative charge (DMPG) correlated with the loss of P0ct helical structure - as seen through the movement of the main spectral minimum towards 200 nm (Fig. 2). NaF had a small unfolding effect on the P0ct secondary structure with all lipid mixtures.

### The kinetics of initial vesicle nucleation

To visualize the underlying kinetics involved in MBP and P0ct binding to lipid vesicles, stopped flow measurements were carried out using an SRCD setup. The original goal was to study the folding kinetics of MBP and P0ct when encountering lipid membranes, but initial measurements and optimization attempts revealed that the proteins gained their folding completely within the dead time of the instrument (∼2 ms), which made the underlying kinetics practically impossible to follow.

Despite the setback with protein folding kinetics, a strong decay in SRCD signal was observed within a 5-10 s measurement window (before the onset of induced photodegradation; data not shown), when proteins were mixed with lipids in the presence of 150 mM NaF or 1 mM CaCl_2_ (Fig. 3A). This resembled exponential decay and could be described with two potentially independent rate constants (Supplementary Table 1). Since we observed similar behaviour in the absence of proteins (Supplementary Fig. S2, Supplementary Table 1), the effect does not stem from changes in protein folding. Considering the wavelength of the used monochromatic beam (195 nm), which is in the size range of large vesicular lipid bodies, and the strength of the decay (spanning tens of mdeg), the effect is caused by light scattering from initial vesicle fusion and/or aggregation, especially since the lipid mixtures and ionic conditions correlate with the observed steady states in turbidimetry (Fig. 1).

**Fig 3.**
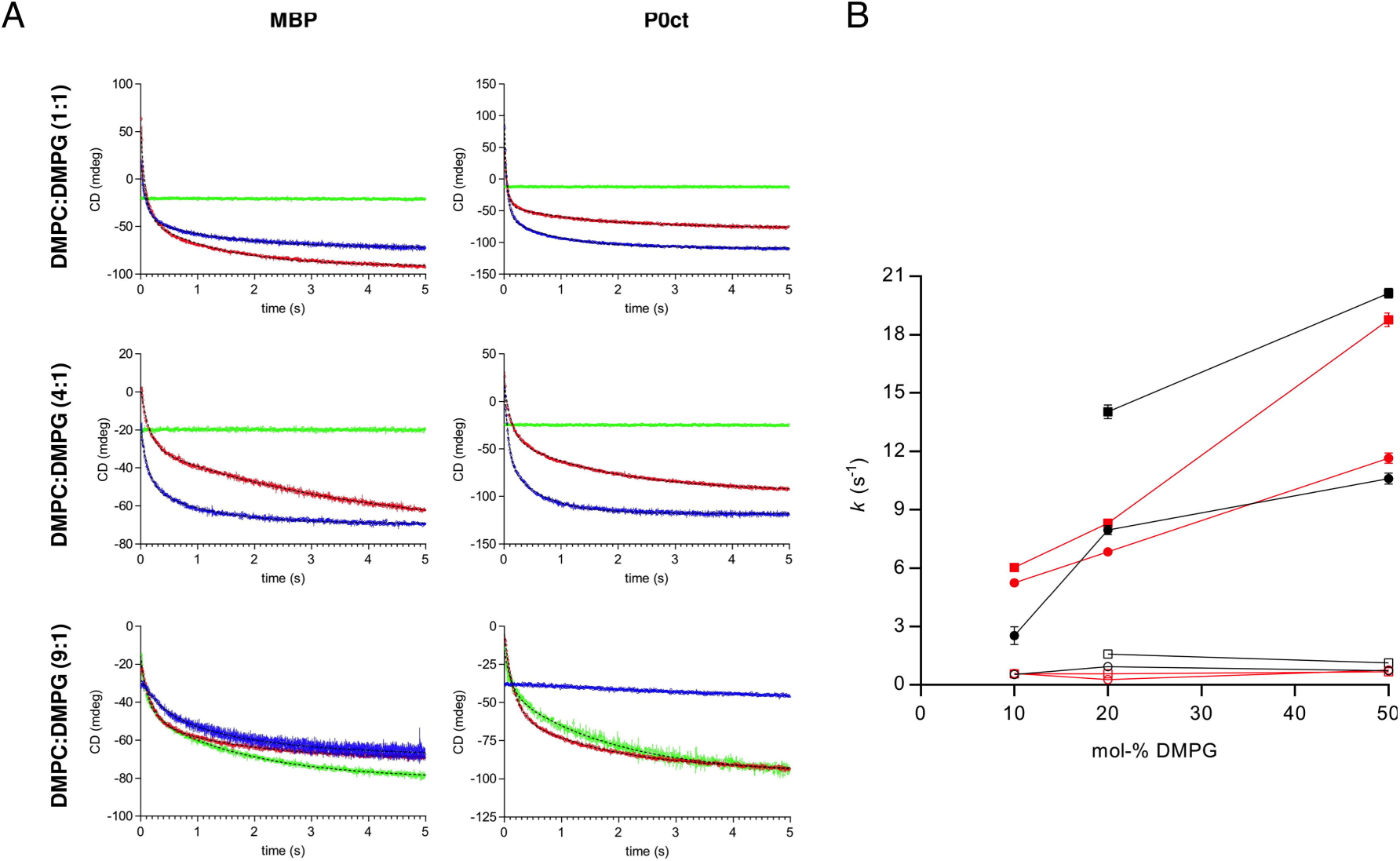
Rapid kinetics of protein-induced lipid turbidity. A. The SRCD signal at 195 nm was monitored for 5 s using stopped-flow measurements. Proteins were mixed with lipids in the absence (green) and presence of 150 mM NaF (blue) and 1 mM CaCl_2_ (red). Error bars represent standard deviation. Fits (dashed black) are shown where data fitting was successful. B. Comparison of kinetic parameters. Evolution of the determined rate constants (*k*_1_, solid markers; *k*_2_, open markers) as a function of DMPC:DMPG molar ratio in 150 mM NaF (black) and 1 mM CaCl_2_ (red). The values for MBP and P0ct are shown as circles and squares, respectively. Error bars represent standard deviation. See Supplementary Table 1 for all rate constants.

While lipids mixed with CaCl_2_ alone produced quite noisy data with individual shots and replicates not overlaying very well, all protein samples had excellent reproducibility both in terms of shot quality and between replicates, which is evident from the small standard deviations. The data could be fitted with confidence, and in most cases, two rate constants were obtained: *k*_1_, which describes a fast event (> 2 s^−1^), and *k*_2_, for a slower event (< 1 s^−1^). The obtained rate constants in the presence of proteins vary between the three different lipid compositions (Fig. 3B). The accuracy of *k*_2_ is rather low, given the limited time window used in the experiments.

In the absence of proteins, the turbidity kinetics induced by Ca^2+^ were similar in DMPC:DMPG (4:1) and (9:1), and clearly slower in DMPC:DMPG (1:1) (Supplementary Fig. S2, Supplementary Table 1), suggesting that electrostatic repulsion plays a role in the observed effect. Considering the fact that the two less charged lipid compositions aggregate with roughly the same rate, the latter effect is more likely to occur due to saturation of the surface with Ca^2+^, although more experiments are required to fully understand the mechanism. It should be noted that the concentration of CaCl_2_ in these experiments was 1 mM, while the total concentration of negative headgroups in the lipid mixtures was either 0.25, 0.1, or 0.05 mM. It appears that the membranes with less charge are fully neutralized, while the neutralization might be partial for the 1:1 mixture.

In protein-lipid samples, in the presence of NaF and CaCl_2_, DMPC:DMPG (1:1) presents a higher *k*_1_ value than the other two lipid compositions. The effects of the two salts on *k*_1_ on turbidity increase with the molar percentage of DMPG (Fig. 3B, Supplementary Table 1).

Interestingly, in DMPC:DMPG (9:1), the aggregation effect was dominated by the 1 mM CaCl_2_ over proteins (Supplementary Table 1); this correlates with the steady-state turbidity under the same condition (Fig. 1C). The rate constants are similar regardless of the presence of protein, indicating that the aggregation behaviour under this condition is governed by the cation. On the other hand, 150 mM NaF inhibits aggregation of DMPC:DMPG (9:1) vesicles (Fig. 1C, Supplementary Table 1), and in salt-free conditions, the proteins display aggregation, which was not the case for the other two vesicle compositions (Fig. 3A). Taken together, the kinetics measurements correlate well with the results from turbidimetry, providing further insights into determinants of membrane binding and stacking.

## Discussion

Myelination requires a delicate balance between myelin protein expression, protein-membrane interactions, and additional regulatory factors, such as calcium (6, 13). Both MBP and P0ct are IDPs in aqueous solution, but upon binding to lipid membranes, they fold into helical structures, at the same time altering the physical properties of the membrane (16, 18, 23, 33, 41). The salt sensitivity of the protein-membrane interactions supports the role of electrostatic interactions in these processes, and the effects of calcium on the molecular interactions are reflective of more specific mechanisms of regulation.

Phospholipid vesicle aggregation was followed using optical density methods as well as SRCD to see how salt content and Ca^2+^ influence the activity and folding of MBP and P0ct. Our results show that while vesicle turbidity increases with ionic strength, both in the steady state and in initial kinetics, the effect of salt varies with membrane surface net charge. Considering the combination of ionic strength and membrane charge, it seems clear that high salt concentrations effectively screen the protein-membrane interactions, while lower concentrations, in fact, have an inducing effect. The results are in line with earlier data on extractability of MBP from myelin and myelin swelling under different conditions (46-48).

The effects of calcium are likely to be more specific than the electrostatic competition caused by near-physiological ionic strength *per se*. Ca^2+^ alone induces spontaneous vesicle turbidity, showing that it affects the surface properties of the membrane. Calcium influences both the steady-state levels and the kinetics of MBP- and P0ct-mediated vesicle aggregation. The binding of divalent cations by MBP has been observed in earlier studies (26, 28, 29), and calcium is known to affect both MBP solubility in purified myelin as well as myelin membrane spacing (48, 49). It, thus, appears that calcium might be able to affect the surface of both the protein and the membrane in a manner that can regulate the direct molecular interaction.

A new SRCD-based time-resolved technique was used to follow the kinetics of vesicle aggregation by MBP and P0ct, as well as by Ca^2+^ alone. While it is at the moment unclear what the rate constants actually represent, some conditions are physiologically relevant, and the observed differences in kinetics may be relevant to myelin membrane stacking. Considering the *k*_1_ values alone, P0ct is faster than MBP at inducing turbidity. The difference could arise from the smaller size of P0ct, which potentially both tumbles in solution faster than MBP and folds/neutralizes more easily upon membrane interaction. Due to its larger size, MBP will experience more steric and repulsive effects from protein molecules already bound to the membrane. The smaller P0ct may possess higher accessibility to the vesicle surface.

Several studies have highlighted calcium as a major regulator of proper myelination (5, 6, 11, 12, 36, 50), and in addition to being crucial for signalling and cellular homeostasis, it has been suggested to directly affect MBP-membrane interactions (14, 36, 48). Phosphoinositides are important constituents of the myelin membrane, involved in MBP-membrane interactions regulated by Ca^2+^(36). Various mechanisms have been proposed for the effects of calcium on myelin structure and formation; these include activation of calpains, possibly degrading MBP, the involvement of calmodulin, signalling pathways, cytoskeletal events, as well as effects on protein-lipid interactions (6, 11–14, 36).

It was proposed that a “sweet spot” concentration of calcium exists during myelination, which promotes normal myelin formation (13). Deviations from this range might be relevant for dys/demyelination. In fact, similar effects have been observed in early studies on MBP extraction from nerve tissue (43, 48). Here, concentrations around the physiological [Ca^2+^] were used to follow myelin protein-membrane interactions, and indeed, concentration dependence around 1 mM [Ca^2+^] is evident. Together with the membrane surface charge, calcium may either promote or inhibit lipid membrane stacking induced by myelin proteins. The obvious next steps would involve studies on the effects of calcium on myelin protein-membrane interactions in more physiological lipid compositions corresponding to the myelin cytoplasmic leaflet; the experiments here using simplified lipid mixtures pave the way for these studies.

MBP and P0 are not related by sequence; however, as shown by us and others, the physicochemical properties of MBP and the P0 cytoplasmic domain are very similar (16, 18, 23, 33, 41, 42). Both bind to membrane surfaces through electrostatic interactions, get embedded and fold, and affect properties of the lipid bilayer. Functional similarity is also true considering the effects of ionic strength and calcium studied here. In addition, other myelin proteins, such as P2 in the PNS and PLP in the CNS (51-54), have likely roles in membrane stacking and will have to work in synergy with MBP and P0 during myelination. Our observations point towards common regulatory mechanisms for the function of different myelin proteins in membrane stacking during myelin maturation and maintenance.

## Acknowledgements

This work was financially supported by the Norwegian Research Council (SYNKNØYT programme). We acknowledge the ASTRID2 synchrotron facility and the Biophysics, Structural Biology, and Screening (BiSS) facility at the University of Bergen. The research leading to this result has been supported by the project CALIPSOplus under the Grant Agreement 730872 from the EU Framework Programme for Research and Innovation HORIZON 2020.

## Author contributions

Study design, original text and figures: A.R., P.K.

Prepared samples and performed experiments: A.R.

Processed and analyzed data: A.R., N.C.J., S.V.H., P.K.

Review & editing: A.R., N.C.J., S.V.H, P.K.

Supervision: P.K.

## Data availability

The datasets generated and analyzed during the current study are available from the corresponding author on reasonable request.

