## Supplementary Information for "Ionic strength and calcium regulate the membrane interactions of myelin basic protein and the cytoplasmic domain of myelin protein zero"

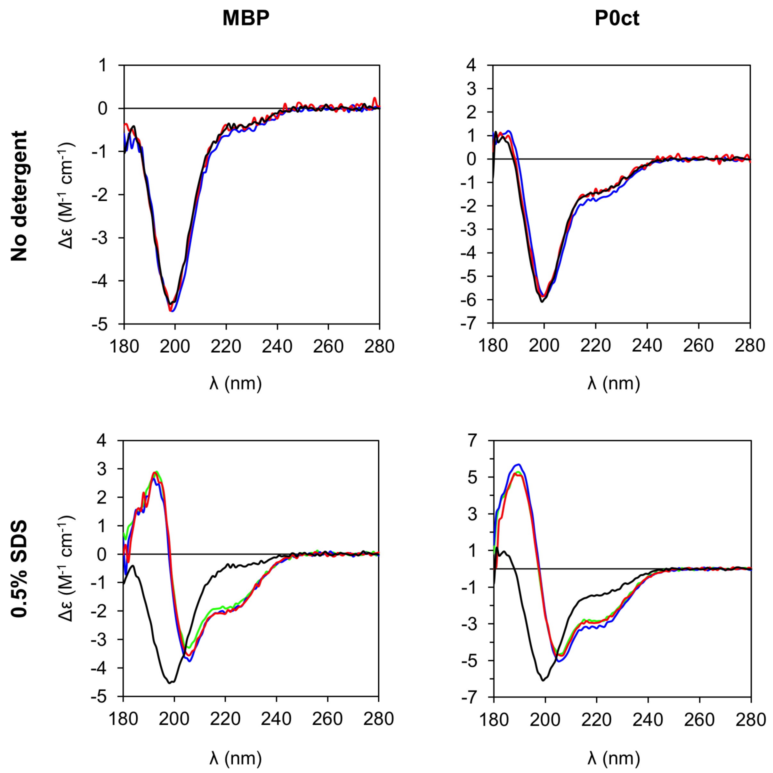


**Supplementary Fig. S1. SRCD control experiments.** The folding of MBP and P0ct in the absence (top) and presence (bottom) of 0.5% SDS as determined by SRCD spectroscopy. No additive (green); 150 mM NaF (blue); 1 mM $CaCl_2$ (red). Protein controls without additives or detergents in water (black) are plotted for reference in each panel.

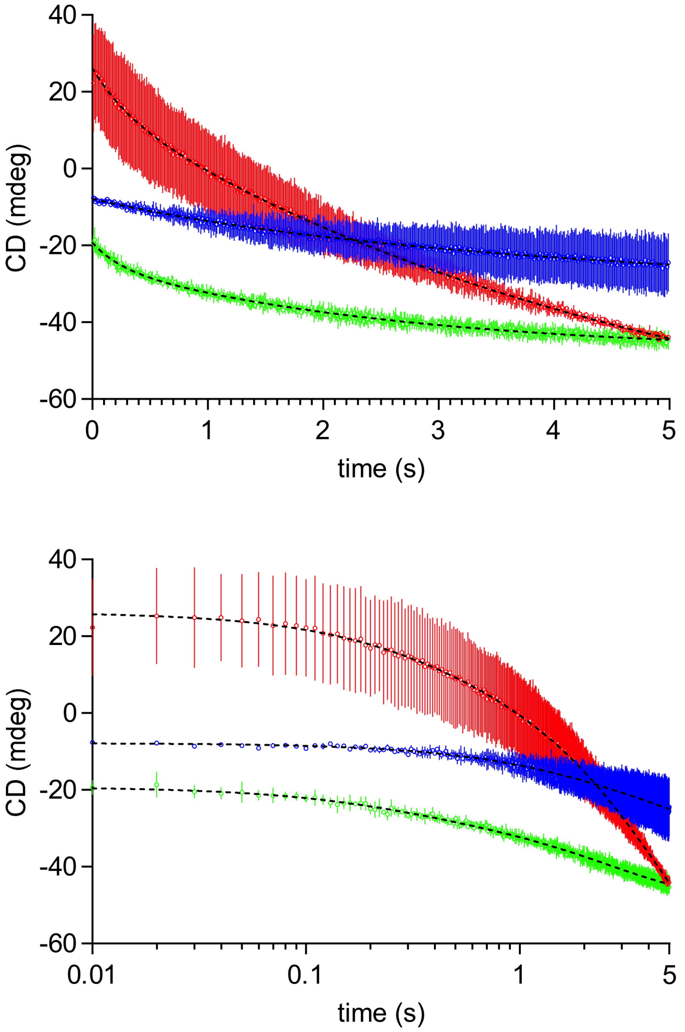


**Supplementary Fig. S2. Rapid kinetics of $Ca^{2+}$-induced initial lipid turbidity.** Evolution of the SRCD signal at 195 nm was monitored for 5 s using stopped flow measurements. For clarity, linear (top) and logarithmic (bottom) time scales are shown. $Ca^{2+}$ was mixed at a final concentration of 1 mM with 2.5 mM DMPC:DMPG 1:1 (green), 4:1 (blue), and 9:1 (red) molar lipid ratios. Error bars represent standard deviation. Fits (dashed black) are shown; see Supplementary Table 1 for rate constants.

**Supplementary Table 1. Two-phase exponential decay rate order parameters from rapid kinetics SRCD experiments.** Values marked with a dash could not be fitted, as the CD signal decayed very little, if at all, within the measurement time window. All errors represent standard deviation.

| Protein | DMPC:DMPG ratio | No additive | | | | 150 mM NaF | | | | 1 mM CaCl2 | | | |
|---|---|---|---|---|---|---|---|---|---|---|---|---|---|
| | | $k_1$ ($s^{-1}$) | $k_2$ ($s^{-1}$) | $k_1/k_2$ | $R^2$ | $k_1$ ($s^{-1}$) | $k_2$ ($s^{-1}$) | $k_1/k_2$ | $R^2$ | $k_1$ ($s^{-1}$) | $k_2$ ($s^{-1}$) | $k_1/k_2$ | $R^2$ |
| No protein | 1:1 | - | - | - | - | - | - | - | - | 0.29 ± 0.04[a] | - | - | 0.6250[a] |
| | 4:1 | - | - | - | - | - | - | - | - | 3.23 ± 1.38 | 0.20 ± 0.03 | 15.89 ± 5.09 | 0.9291 |
| | 9:1 | - | - | - | - | - | - | - | - | 4.20 ± 0.78 | 0.39 ± 0.03 | 10.89 ± 1.56 | 0.9564 |
| MBP | 1:1 | - | - | - | - | 10.59 ± 0.29 | 0.73 ± 0.02 | 14.55 ± 0.37 | 0.9800 | 11.65 ± 0.26[b] | 0.76 ± 0.01[b] | 15.23 ± 0.30 | 0.9878[b] |
| | 4:1 | - | - | - | - | 7.95 ± 0.24 | 0.93 ± 0.02 | 8.59 ± 0.20 | 0.9868 | 6.84 ± 0.17 | 0.27 ± 0.01 | 25.34 ± 0.62 | 0.9934 |
| | 9:1 | 8.52 ± 0.25 | 0.56 ± 0.01 | 15.13 ± 0.36 | 0.9927 | 2.53 ± 0.46 | 0.54 ± 0.09 | 4.68 ± 0.41 | 0.9376 | 5.24 ± 0.11 | 0.58 ± 0.02 | 9.04 ± 0.18 | 0.9924 |
| P0ct | 1:1 | - | - | - | - | 20.14 ± 0.25 | 1.12 ± 0.01 | 17.96 ± 0.22 | 0.9934 | 18.77 ± 0.36 | 0.67 ± 0.01 | 27.95 ± 0.61 | 0.9837 |
| | 4:1 | - | - | - | - | 14.03 ± 0.36 | 1.58 ± 0.03 | 8.89 ± 0.18 | 0.9893 | 8.30 ± 0.16 | 0.57 ± 0.01 | 14.50 ± 0.23 | 0.9954 |
| | 9:1 | 7.79 ± 0.61 | 0.53 ± 0.01 | 14.59 ± 0.97 | 0.9785 | - | - | - | - | 6.03 ± 0.09 | 0.57 ± 0.01 | 10.57 ± 0.13 | 0.9963 |

[a] Data fit best into one-phase exponential decay function, but the change in SRCD signal overall is very small.

[b] Dataset is visibly more complicated that what can be described with two rate constants.
